# Transcriptional and cellular signatures of cortical morphometric similarity remodelling in chronic pain

**DOI:** 10.1101/2021.03.24.436777

**Authors:** Daniel Martins, Ottavia Dipasquale, Mattia Veronese, Federico Turkheimer, Marco L. Loggia, Stephen McMahon, Matthew A Howard, Steven CR Williams

**Affiliations:** Department of Neuroimaging, Institute of Psychiatry, Psychology and Neuroscience, King’s College London, De Crespigny Park, London SE5 8AF, UK; Athinoula A. Martinos Center for Biomedical Imaging, Department of Radiology, Massachusetts General Hospital and Harvard Medical School; Wolfson CARD, Institute of Psychiatry, Psychology & Neuroscience, King’s College London, London SE1 1UL

**Keywords:** Chronic pain, morphometric similarity, neuroinflammation, Allen Brain Atlas, transcriptomics

## Abstract

Chronic pain is a highly debilitating and poorly understood condition. Here, we attempt to advance our understanding of the brain mechanisms driving chronic pain by investigating alterations in morphometric similarity (MS) and corresponding transcriptomic and cellular signatures, in three cohorts of patients with distinct chronic pain syndromes (knee osteoarthritis, low back pain and fibromyalgia). We uncover a novel pattern of cortical MS remodelling involving mostly MS increases in the insula and limbic cortex, which cuts across the boundaries of specific pain syndromes. We show that cortical MS remodelling in chronic pain spatially correlates with the brain-wide expression of genes involved in the glial immune response and neuronal plasticity. Cortical remodelling in chronic pain might involve a disruption of multiple elements of the cellular architecture of the brain. Therefore, multi-target therapeutic approaches tackling both glial activation and neuronal hyperexcitability might better encompass the full neurobiology of chronic pain.

## Introduction

Chronic pain is a highly debilitating condition^1-3^ characterized by persistence of pain beyond normal healing time^1^. Although the number of available treatments has grown substantially (i.e. antidepressants, anticonvulsants, opioids)^4^ over the last decades, treatment response to current standard treatments in patients with chronic pain is overall poor^5^. This problem is further amplified by the fact that most of the available pharmacological treatments are accompanied by considerable side effects and risk of misuse (i.e. opioids)^6^, which motivates high rates of treatment non-adherence^7^. Identifying new potential targets for drug development in chronic pain has become over the years a priority for researchers and clinicians. However, so far, therapeutic advances have been limited^8^.

To a large extent, this lack of progress in the development of new therapeutics stems from a poor understanding of the mechanisms underlying chronic pain. Currently, it is generally accepted that chronic pain is a multifactorial entity entailing physical, psychological, emotional, and social aspects^1^. Over the years, chronic pain has been increasingly recognized as a disorder of the central nervous system, including the brain^9^. Animal studies have offered insights into key central mechanisms that might contribute to chronic pain, such as the sensitization of an array of nervous system pathways, imbalances in the facilitatory and inhibitory descending modulation pathways from the brain that regulate the transmission of noxious information in the spinal cord, neuroinflammation and glial dysfunction, among others^10-14^. These findings have fuelled a number of neuroimaging studies in humans seeking to unravel how chronic pain affects brain structure, functioning and neurochemistry, and what form of brain pathophysiology in these patients might be amenable to be targeted by different pharmacological and non-pharmacological treatments^15,16^. Using different neuroimaging techniques, a rich body of evidence has provided evidence that the majority of chronic pain syndromes is associated with a number of spatially-patterned structural and functional alterations in different cortical (such as the insular, somatosensory, motor and association cortices), subcortical regions (including the thalamus, basal ganglia, hippocampus and the amygdala), and in the brainstem (including the midbrain, rostral ventromedial medulla and the periaqueductal gray)^9,17-19^. A parsimonious explanation for this distributed pattern of brain changes is that it might reflect disruption of large-scale brain networks comprising anatomically connected brain areas^20^. This hypothesis has received empirical support from a number of studies showing aberrant functional^21-24^ and structural^25-28^ configuration of the brain network topology in patients with different chronic pain syndromes. However, testing this connectivity hypothesis has been constrained by challenges in measuring anatomical connectivity in humans^29^. For instance, the principal imaging methods available for this purpose are diffusion weighted imaging (DWI) tractography and structural covariance analysis of structural MRI. The use of DWI-based tractography to estimate the connectivity strength of long distance projections, such as those between hemispheres, remains challenging^30^. On the other hand, structural covariance analysis is not applicable at the single-subject level, and since it relies on group-based covariance, it requires large sample sizes to be reliably estimated^29^.

Morphometric similarity (MS) mapping has recently emerged as a new approach to construct whole-brain anatomical networks for individual subjects, overcoming some of the methodological limitations highlighted above^31-33^. It quantifies the similarity between cortical regions in terms of multiple MRI parameters measured in each area^33^. This metric has close associations with the cytoarchitectonic properties of the cortex^33^. In keeping with histological results indicating that cytoarchitectonically similar areas of the cortex are more likely to be anatomically connected, MS in the macaque cortex was correlated with tract-tracing measurements of axonal connectivity^33^. MS mapping is a reliable and robust method and has been demonstrated to capture inter-individual differences in cognition^33^ and clinical abnormalities in psychosis^31^, neurogenetic disorders^32^ and major depressive disorder (MDD)^34^. Nevertheless, its use for uncovering morphometric differences in the brain of patients with chronic pain syndromes has never been attempted before.

One aspect of MS mapping that makes it particularly attractive to study morphometric changes in the brain of patients with chronic pain and potentially advance our understanding of the neurobiological mechanisms underlying these changes is its tight relationship with gene expression. For instance, regions with strong shared patterns of MS also present high levels of gene co-expression^33^. Furthermore, changes in regional MS have been demonstrated to be able to uncover transcriptomic and cellular profiles of regional brain vulnerability to brain disorders^31,32,34^. Indeed, while previous studies have shown that 16-50% variation in chronic pain development is heritable^35,36^, how disease-related alterations at the microscopic genetic architecture drive macroscopic brain abnormalities in chronic pain syndromes is currently largely unknown.

Here, we sought to bridge these gaps by investigating changes in MS in three independent case-control studies of patients with chronic pain syndromes: knee osteoarthritis (OA), chronic low back pain (CLBP) and fibromyalgia (FM). We mapped case–control morphometric similarity differences at global and regional levels of resolution individually in each dataset, and we tested for differences in MS organization that were consistent across conditions. We also used data from the Allen Human Brain Atlas (AHBA) to test the hypothesis that this MS brain phenotype of chronic pain is correlated with anatomically patterned gene expression (see Figure 1 for an overview of our analysis pipeline). This analytical approach combining imaging and genomic data has been empirically validated^37^ and applied in the context of several neuropsychiatric disorders^31,32,38-40^. We used it to explore among potential molecular and cellular pathways that might underlie MS changes during chronic pain in the hope it might unravel novel aspects of the pathophysiology of chronic pain, which could guide future endeavours in drug development and/or repurposing.

**Figure 1.**
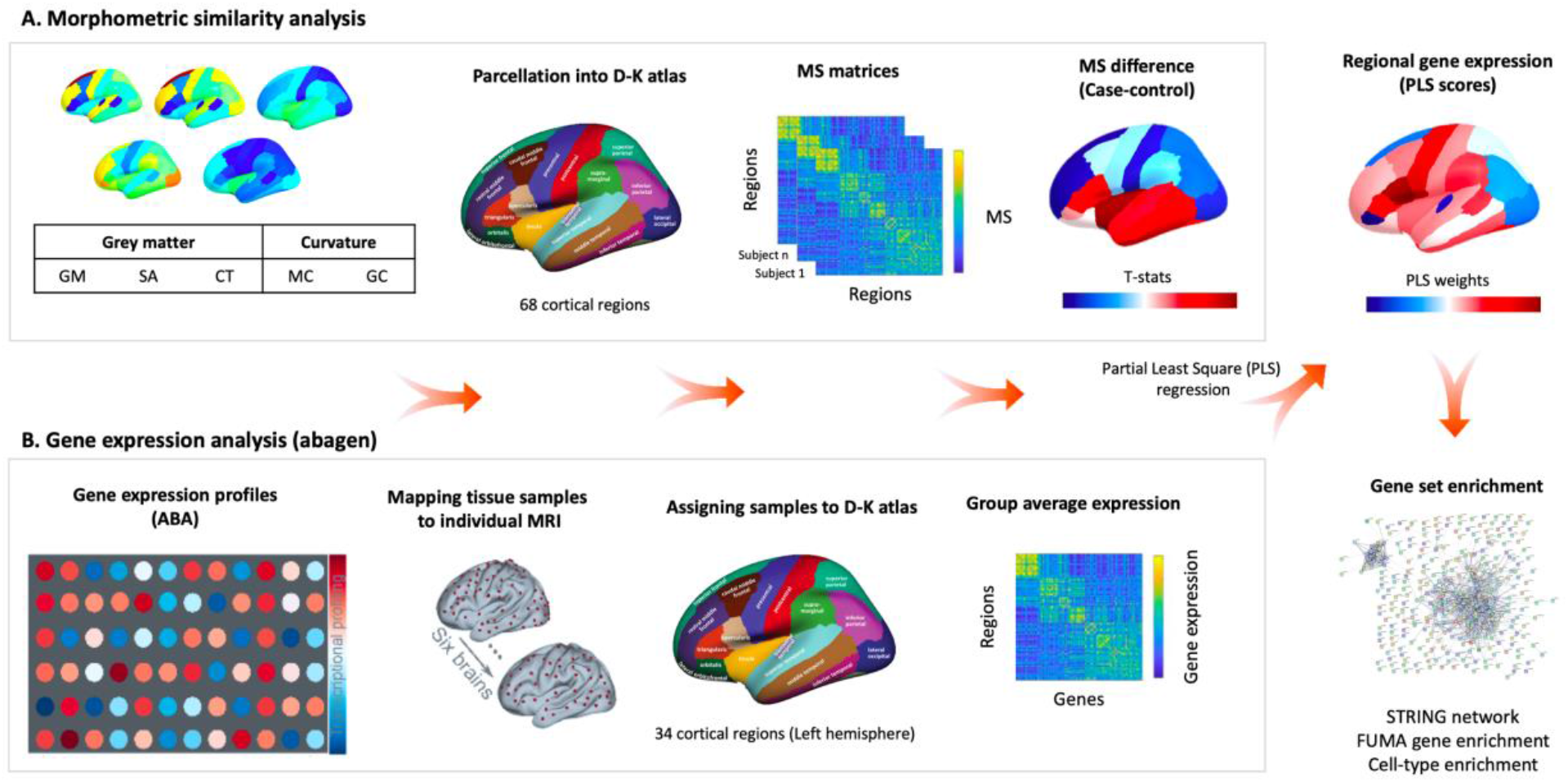
Overview of the analysis pipeline. **(A)** Morphometric similarity (MS) analysis. We constructed individual cortical MS matrices using five structural magnetic resonance imaging features (e.g. grey matter volume (GM), surface area (SA), cortical thickness (CT), mean curvature (MC) and Gaussian curvature (GC)) extracted from 68 cortical regions of the *Desikan-Killiany (DK)* atlas. For each individual, we produced a 68 × 68 MS matrix by correlating the normalized (z-scores) values of the five structural features between each pair of regions in the atlas. For each region, we averaged across all the edges involving that area to obtain a singular representation of the mean MS score for that region. We then computed case-control differences for each region, while accounting for age, gender and total intracranial volume. **(B)** Gene expression analysis. We used *abagen* to obtain gene expression profiles from the Allen Human Brain Atlas (AHBA) in 68 regions (left hemisphere only) across the six post-mortem brains sampled in this atlas. We excluded all genes with normalized expression values below the background (15,633 genes met this criterion). When more than one probe was available for a certain gene, we selected the probe with higher consistency in expression across the 6 donors. We used partial least squares regression (PLS) to rank all genes according to their association with the case-control changes in MS. Finally, we performed a set of enrichment analyses on the top genes positively or negatively associated with case-control differences in MS.

## Results

### Morphometric similarity is a reproducible measure in healthy controls across independent studies

The cortical maps of regional MS in Supplementary Figure S1 summarizes the anatomical distribution of areas of positive and negative MS in healthy controls from each of the three datasets. The patterns of regional distribution of MS were highly correlated across healthy controls from the three datasets (Supplementary Figure S2). The results are similar to those reported in other independent samples^31-33^, with positive MS in frontal and temporal cortical areas (indicating high levels of similarity with the rest of the cortex) and negative MS in occipital and cingulate cortices (indicating low levels of similarity with the rest of the cortex; hence, high levels of differentiation from the rest of the cortex). This confirms the replicability of this pattern of regional MS in healthy individuals.

### Chronic pain patients do not differ from healthy controls in global morphometric similarity

Regional MS had an approximately normal distribution over all 68 cortical regions (after regressing age, sex, and intracranial volume) in both patients and healthy controls from all three datasets (Figure 2 – upper panel). We did not find any significant case-control differences in global MS in any of the three datasets (OA: T(75)=0.98, p_unc_ = 0.33; CLBP: T(61) = -0.56, p_unc_ = 0.58, FM: T(39) = -0.84, p_unc_ = 0.41) (Figure 2 – lower panel).

**Figure 2.**
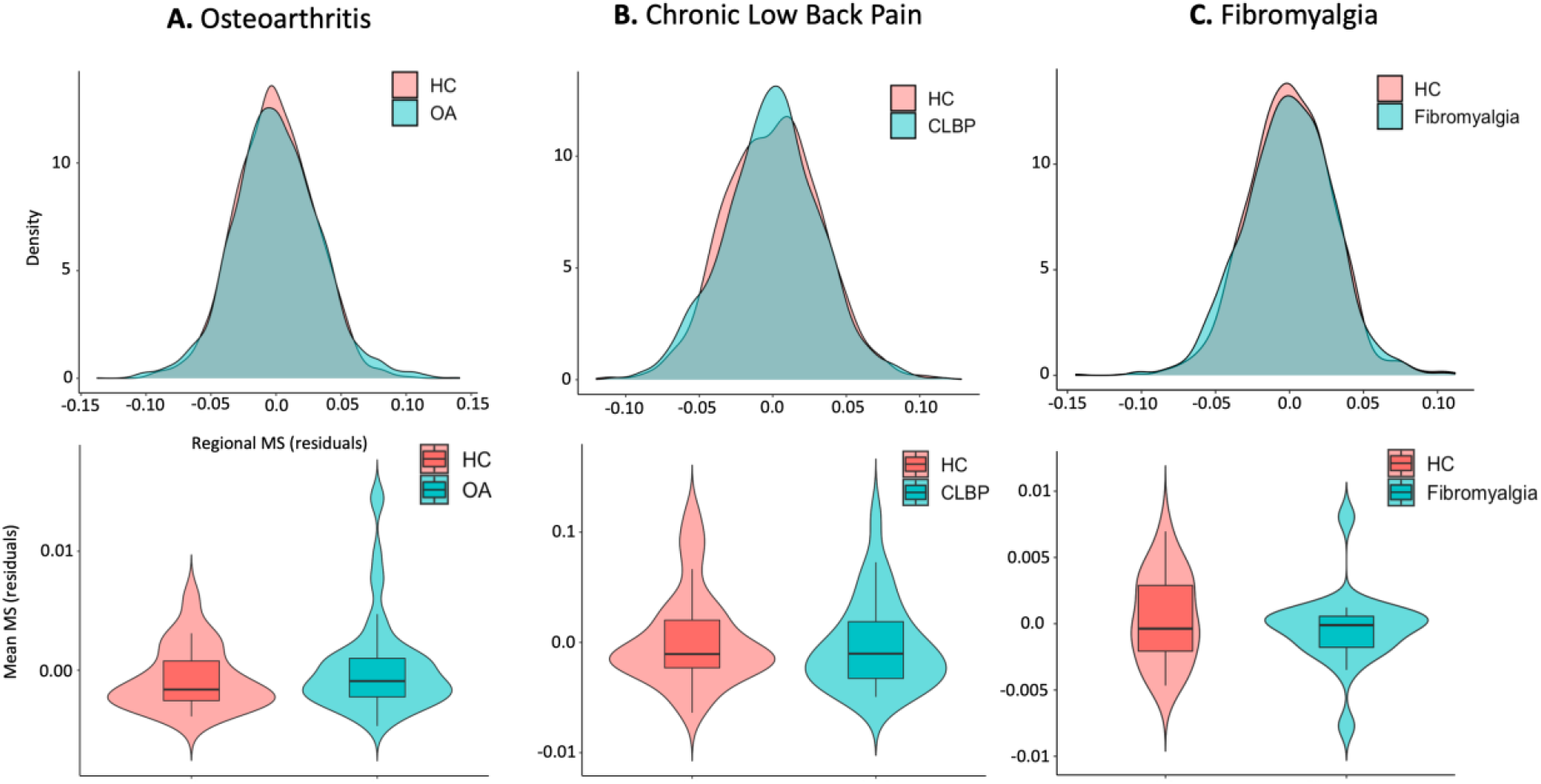
Case-control differences in global morphometric similarity. In the upper panel, we present case and control distributions of the residual regional morphometric similarity (MS) strength (i.e., the average similarity of each region to all other regions) after regressing out the effects of age, gender and intracranial volume, for each dataset separately. In the lower panel, we present case-control comparisons of the global MS. To calculate global MS, we averaged the residual regional MS strength across all regions for each subject. *Abbreviations*: OA – Osteoarthritis; CLBP – Chronic Low Back Pain; HC – Healthy Controls

### Differences in regional morphometric similarity in patients with chronic pain syndromes as compared to healthy controls

The cortical maps in Figure 3 – upper panel summarize the distribution of case-control changes in cortical MS for each chronic pain condition. In Figure 3 – lower panel, we highlight only regions with case-control differences significant at p < 0.05, uncorrected (none of these regions survived FDR correction). In the OA dataset, we found decreases in MS in the left superior frontal gyrus, right pericalcarine cortex and in the left posterior cingulate, and increases in the left insula and inferior temporal gyrus, and in the right bank of the superior temporal sulcus and right inferior temporal gyrus. In the CLBP dataset, we found decreases in MS in the left and right superior parietal gyri, and left lateral occipital cortex; and increases in the left entorhinal cortex and caudal anterior cingulate, and in the right insula. In the fibromyalgia dataset, we found decreases in the left superior parietal, medial and inferior temporal and fusiform gyrus; and increases in the left and right isthmus of the cingulate, left posterior cingulate, and entorhinal and parahippocampal cortices. Changes in regional MS correlated positively between conditions (Supplementary Figure S3), suggesting the existence of a shared pattern of regional MS changes across the three conditions. To further support the existence of this pattern, we performed a principal component analysis (PCA) on the regional MS changes of the three conditions, finding that the first PC explained the majority of variance (64.45%) in case-control changes across the three conditions (PC1).

**Figure 3.**
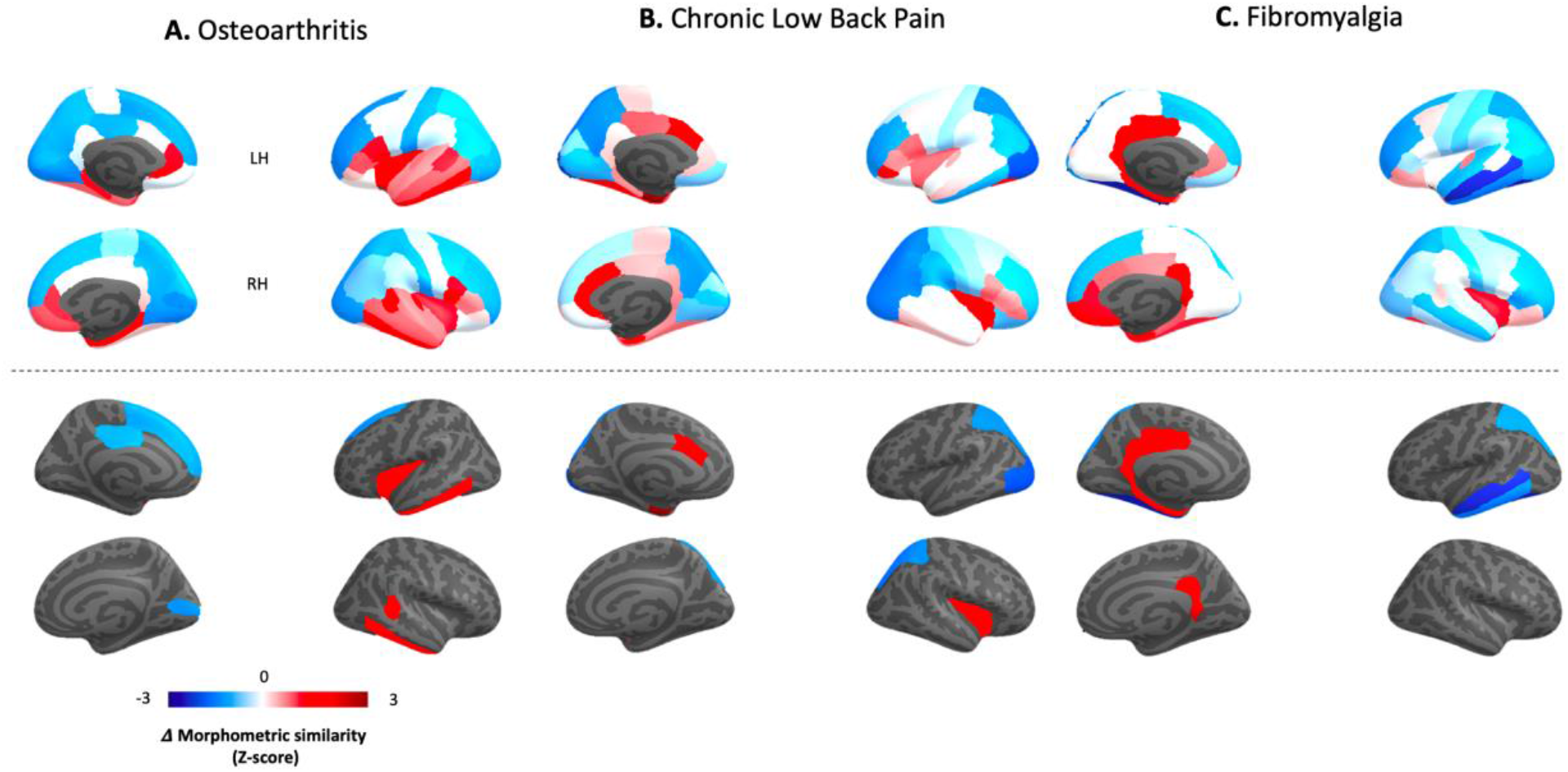
Case-control differences in regional morphometric similarity. In this figure, we present cortical maps of the distribution of Z-scores quantifying case–control differences in regional morphometric similarity (Δ Morphometric similarity) for each condition. In the upper panel, we present unthresholded maps. In the lower panel, we present only regions where we found case-control differences for a threshold of p<0.05, uncorrected. Note that none of these regions survived FDR correction for the total number of regions tested within each condition.

Since we used five different cortical features to estimate MS (gray matter volume, surface area, cortical thickness, gaussian curvature and mean curvature), we tested for differential contributions of single cortical features to the observed regional MS changes by recomputing the condition-specific MS change maps excluding each individual cortical feature at a time prior to MS calculation. We then determined which of these single-feature exclusions most changed the topography of the observed MS changes by identifying the leave-one-feature-out map that correlated the least with the total map. This leave-one-out procedure showed that cortical thickness was the feature that most contributed to the topography of observed regional MS changes across the three conditions (Supplementary Figures S4 and S5). Yet, all leave-one-out maps were positively correlated in all of the three conditions (Supplementary Figures S4).

Chronic pain is often comorbid with major depressive disorder (MDD)^41^. Several preclinical and clinical studies have found considerable overlaps between chronic pain- and depression-induced neuroplasticity^42^. To investigate whether the regional morphometric similarity changes we report here might predominantly reflect neuroplasticity associated with mood alterations other than chronic pain per se, we used another openly available dataset including high resolution structural brain data from unmedicated patients with major depressive disorder and healthy controls^43^. We used these data to define the pattern of changes in regional MS associated with MDD and to investigate whether changes in regional MS in the MDD dataset can predict those we observed during chronic pain. We found a considerably different pattern of MS changes in unmedicated MDD patients, as compared to never depressed healthy controls (Supplementary Figure S6A). Morphometric similarity changes in the MDD dataset did not correlate with MS changes in OA and correlated negatively with MS changes in CLBP and fibromyalgia. We also found a negative correlation between MS changes in MDD and PC1 capturing the cross-condition pattern of changes (Supplementary Figure S6B). Hence, the pattern of MS changes we report here for chronic pain patients is unlikely to simply reflect neuroplastic changes associated with comorbid MDD.

### Mapping case-control differences in regional morphometric similarity to established patterns of cytoarchitectonic cortical organization

To help us to contextualize the case-control differences in regional MS we observed for the different chronic pain conditions, we mapped them onto well-established patterns of cytoarchitectonic organization of the cortex as defined by the *von Economo* atlas of the cortex^**37**^. In OA, we found an increase in MS in the insular cortex (p < 0.05, uncorrected). In both CLBP and fibromyalgia, we found increases in MS in the limbic cortex (p < 0.05, uncorrected). We also found decreases in association cortex A in fibromyalgia (p < 0.05, uncorrected) (Figure 4B and Supplementary Table S1). None of these changes survived FDR correction.

**Figure 4.**
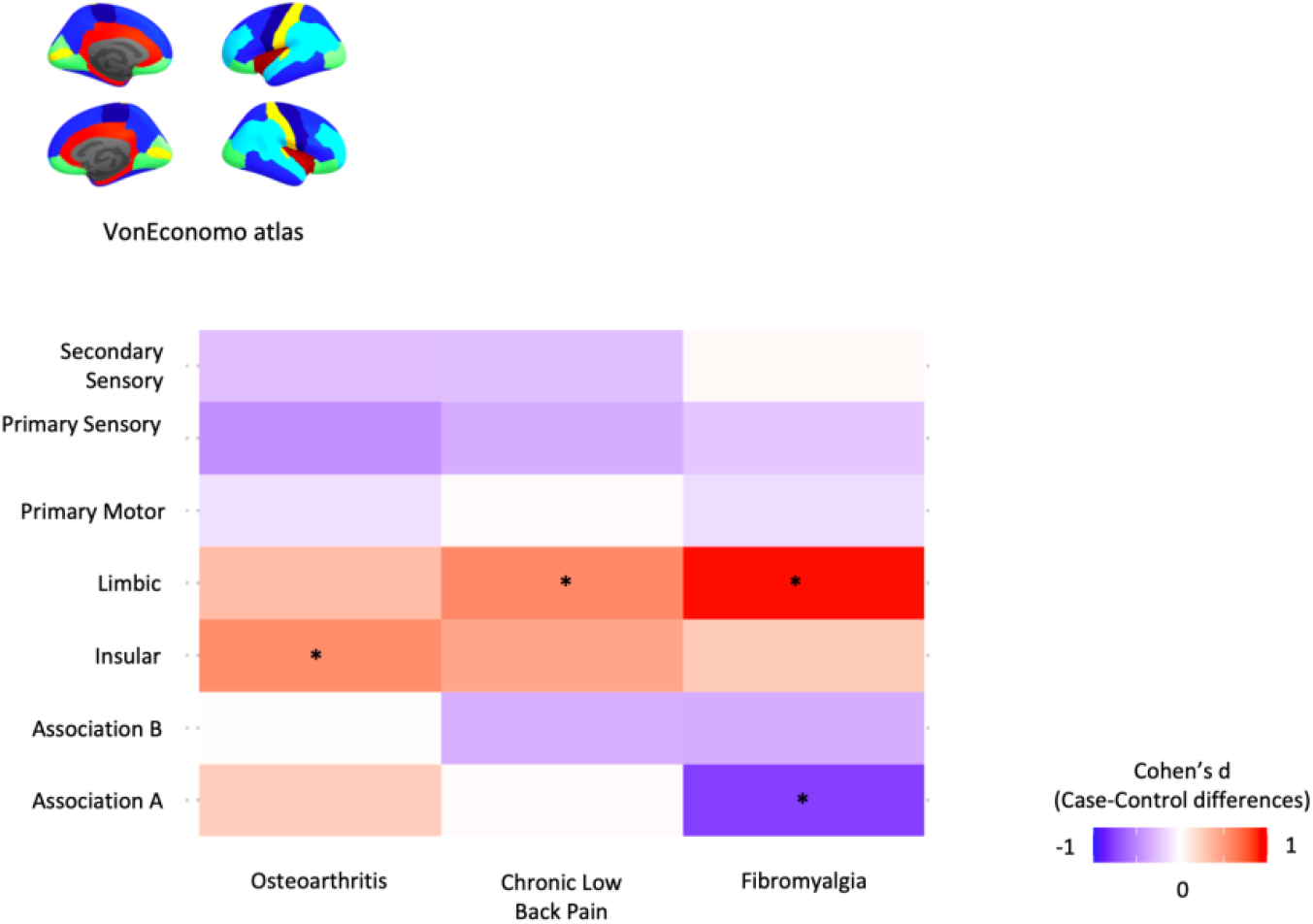
Mapping case-control differences in regional morphometric similarity to established patterns of cytoarchitectonic cortical organization. To help us to contextualize the case-control differences in regional morphometric similarity we observed for the different chronic pain conditions, we mapped them in relation to the von Economo atlas of cortex classified by cytoarchitectonic criteria. For each subject, we quantified MS within each parcel of these atlas (after regressing out age, sex and intracranial volume) and then performed case-control comparisons using independent sample-T tests. The colours in the tile plots represent the Cohen’s d effect size of case-control differences for each condition. * highlights significant case-control differences, for p < 0.05, uncorrected.

### “Hub susceptibility”: Associations between case-control differences in regional morphometric similarity and regional morphometric similarity in healthy controls

Previous studies have shown that highly connected “hub” regions are the most likely to show reduced connectivity in disease in fMRI and diffusion tensor imaging (DTI) brain networks^44^. Hence, here we also tested this “hub susceptibility” model by investigating relationships between case-control changes in regional MS and the typical pattern of regional MS distribution in healthy controls. Across the three datasets, we found significant negative correlations between these two variables (OA – Spearman Rho = -0.397, p_spin_=0.001; CLBP - Spearman Rho = -0.379, p_spin_=0.003; FM - Spearman Rho = -0.522, p_spin_=1.7×10^−04^) (Figure 5 – upper panel). Moreover, we found that most regional increases map to regions typically showing low MS in controls, while most regional decreases map to regions of high MS in controls (Figure 5 – lower panel). These findings suggest that chronic pain is associated with decoupling of MS “hubs” and de-differentiation of highly differentiated regions.

**Figure 5.**
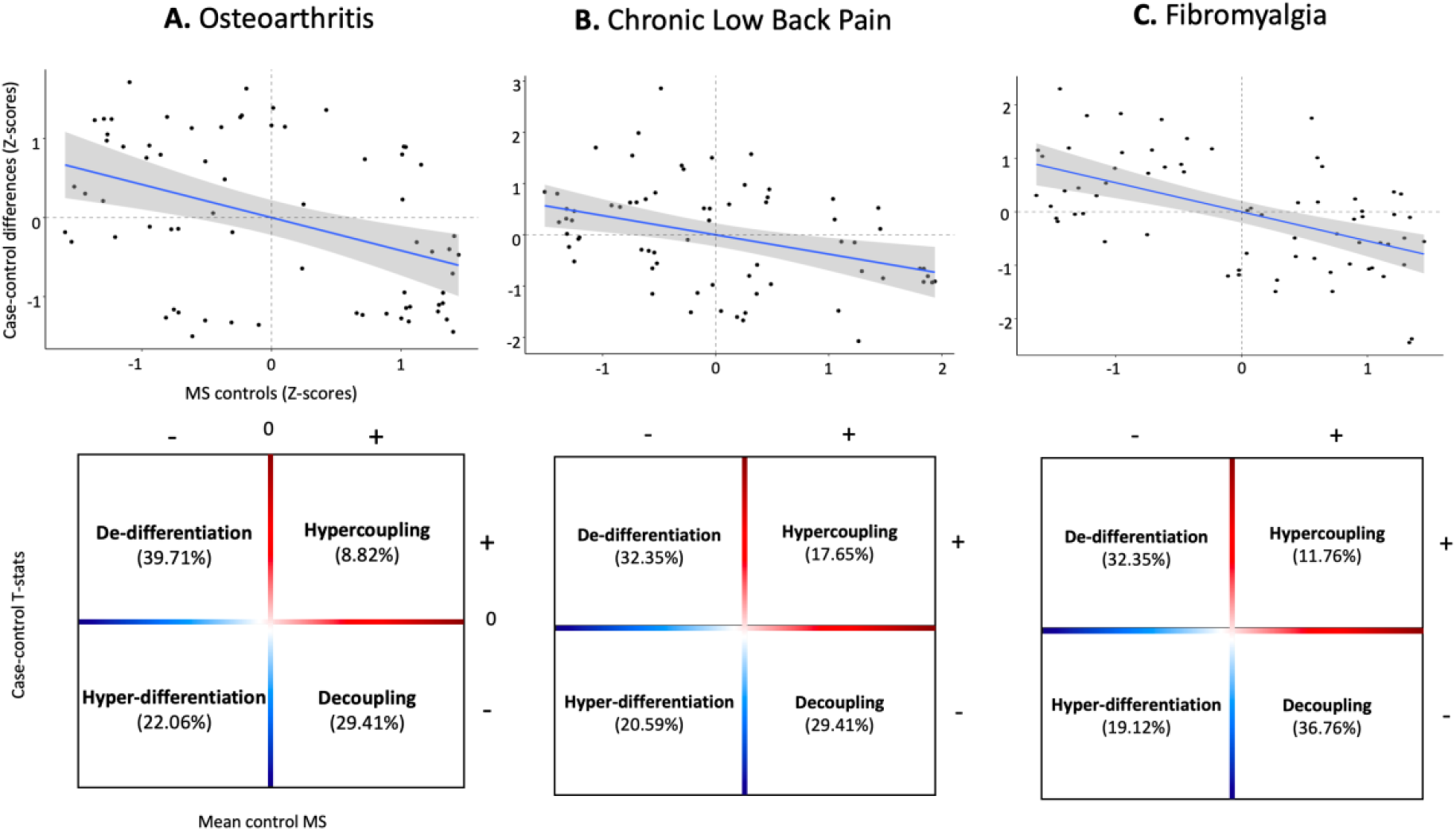
Chronic pain is associated with decoupling of morphometric similarity “hubs” and de-differentiation of highly differentiated regions. In the upper panel, we present scatterplots depicting significant negative associations between case-control differences in regional morphometric similarity and the mean morphometric similarity of each region in healthy controls. In the lower panel, we subdivided these scatter plots in four quadrants according to the relationship between the distribution of case-control differences and mean regional morphometric similarity in healthy controls. The upper left quadrant includes regions of low MS in healthy controls (highly differentiated from the rest of the cortex) that increase their MS during with the rest of the cortex during chronic pain (De-differentiation). The upper right quadrant includes regions of high MS in healthy controls (highly connected with the rest of the cortex) that increase their MS during chronic pain (Hypercoupling). The lower left quadrant includes regions of low MS in healthy controls that decrease their MS during chronic pain (Hyper-differentiation). The lower right quadrant includes region of high MS in healthy controls that decrease their MS during chronic pain (Decoupling). Across chronic pain conditions, we observed a predominant pattern of increases in regional MS in regions of low MS (De-differentiation) and decreases in regions of high MS (Decoupling) in healthy controls. Significance was assessed with spatial permutation testing.

### Transcriptomic correlates of cortical morphometric similarity remodelling during chronic pain

We investigated associations between cross-condition changes in MS during chronic pain (PC1) and brain gene expression using partial least square regression (PLS). The first PLS component (PLS1) explained the highest proportion of MS changes (24.4%) and did so above chance (p_boot_ = 0.003). PLS1 gene expression weights were positively correlated with cross-condition changes in regional MS (r = 0.494, p_spin_ = 0.013) (Figure 6, panel A). This positive correlation means that genes positively weighted on PLS1 are highly expressed in regions where MS was increased, while negatively weighted genes are highly expressed in regions where MS was decreased in patients.

**Figure 6.**
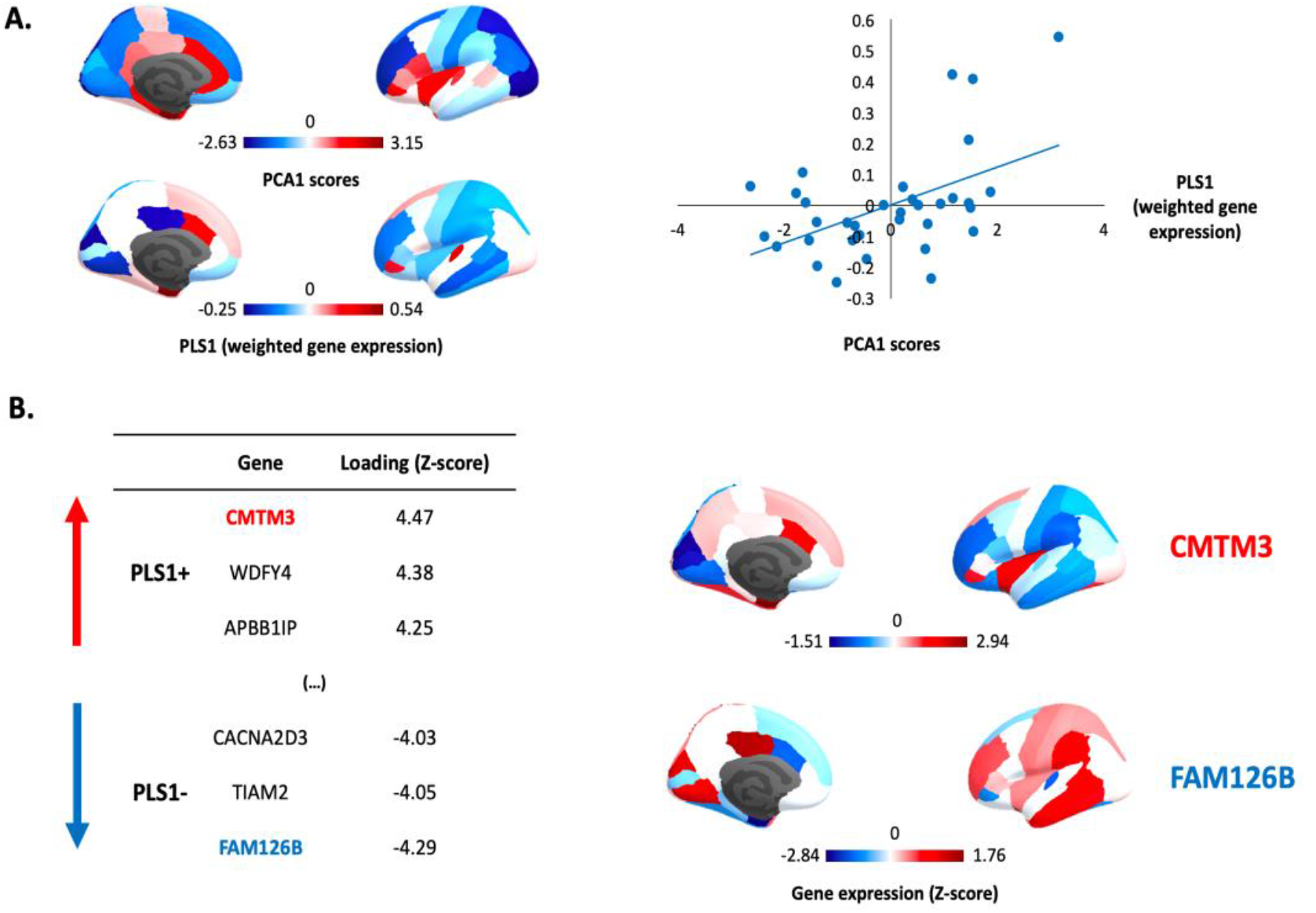
Gene expression profiles related to cross-condition changes in regional morphometric similarity during chronic pain. (A) We summarized case-control changes in regional morphometric similarity (MS) across chronic pain conditions using principal component analysis (PCA). The first component (PC1) explained the large majority of variance 64.45% and was the only component with an eigenvalue > 1. The cortical distribution of the scores of PC1 is depicted in the upper part of panel A. In the lower part, we show the cortical distribution of PLS1 scores summarizing the regional weighted expression of genes associated with cross-condition changes in regional MS during chronic pain. In the scatter plot on the right, we depict a significant positive correlation between PLS1 gene expression weights and the PC1 scores summarizing case–control regional MS differences across conditions. In panel B, we present the top 3 genes positively and negatively associated with PCA1, ranked by the respective loading into PLS1. Loading (Z-score) refers to the weight of each gene in PLS1. Genes with positive weights are highly expressed in regions where MS increases in patients, while genes with negative weights are highly expressed in regions where MS decreases. In the right part of panel B, we provide cortical maps summarizing the regional distribution of the top genes with the highest (CMTM3) or lowest (FAM126B) weights in PLS1.

We found 338 genes with PLS1 weights Z > 3 (which we denoted PLS1+) and 236 genes with Z < -3 (PLS1-) (all p_FDR_ < 0.05). The gene with the highest positive weight was the “Chemokine-Like Factor Superfamily Member 3” (CMTM3), a microglia gene related to immune cytokine activity. The gene with the lowest negative weight was the “Family with Sequence Similarity 126, Member B” (FAM126B), which is part of a complex required to localize phosphatidylinositol 4-kinase to the plasma membrane (Figure 6B).

### Protein-Protein networks and gene-set enrichment

We mapped the network of known interactions between proteins coded by the PLS1+ and PLS1− gene sets (Supplementary Figure S7). For PLS1+, the resulting network had 303 nodes and 615 edges, more than the 239 edges expected by chance (PPI enrichment p-value < 1×10^−16^). Using gene set enrichment analysis, we found enrichment for a number of gene ontology terms – biological pathways broadly mapping to the neuroimmune response axis (Figure 7A). For PLS1-, the resulting network had 225 nodes and 195 edges, more than the 152 edges expected by chance (PPI enrichment p-value 0.0005). Using gene set enrichment analysis, we did not find enrichment for any gene ontology terms, but found four terms from the *Kyoto Encyclopedia of Genes and Genomes* (KEGG) pathways that reached significance. Those were “Calcium signalling”, “Long-term potentiation”, “Taste transduction” and “Type II diabetes mellitus” (Figure 7A).

**Figure 7.**
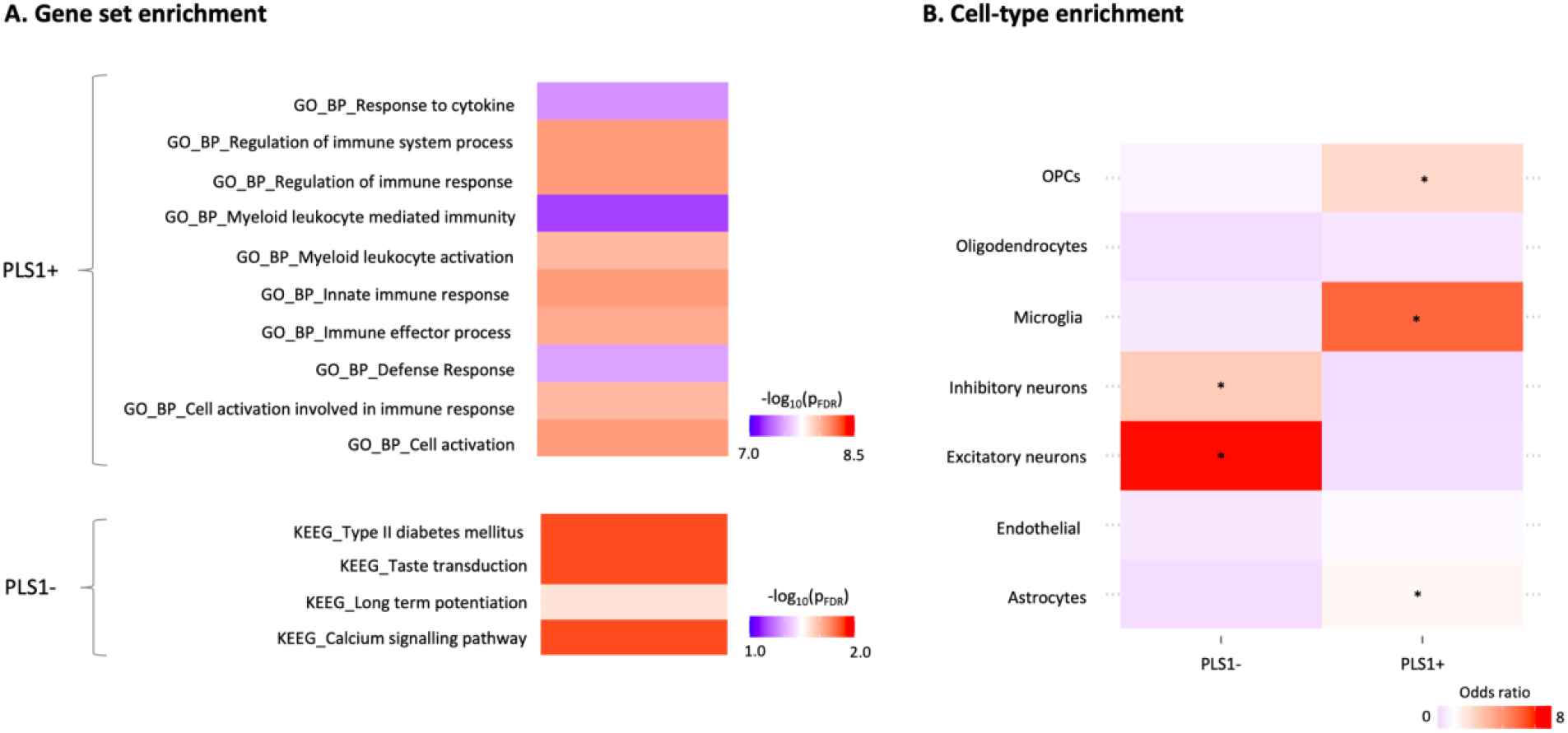
Gene set and cell-type enrichment analyses of the top genes associated with cross-condition changes in regional morphometric similarity during chronic pain. In panel A, we present the results of gene set enrichment analyses on the top genes positively (PLS1+) and negatively (PLS1-) associated with cross-condition changes in regional morphometric similarity during chronic pain, as implemented in the *Functional Mapping and Annotation of Genome-Wide Association Studies* (FUMA) platform. For PLS1+, we present the top 10 gene ontology (GO) – biological pathways terms for which we found significant enrichment, ranked by p-value after FDR correction. For PLS1-, none of the GO terms survived correction, but we found significant enrichment for four terms from the Kyoto Encyclopedia of Genes and Genomes (KEGG) pathways. Colour scale indicates –log(P_FDR_). The full results of the FUMA analysis can be found in supplementary data. In panel B, we present the results of a cell-type enrichment analysis, where we investigated whether PLS1+ and PLS1-include overrepresentation of genes typically expressed in specific brain cell-types. Enrichment was quantified using odds ratio (OR) and significance calculated with a Fisher’s exact test. Colour scale indicates OR. Higher ORs (in red) indicate higher enrichment in genes of a certain cell class. The asterisk denotes cell classes for which we found significant enrichment, after correcting for the number of cell types tested (*p_FDR_ < 0.05). *Abbreviations*: FDR – False Discovery Rate; OPCs – Oligodendrocyte precursor cells.

### Enrichment for transcriptional signatures of canonical brain cell-types

We also performed cell-type enrichment analysis using omnibus lists of gene expression in different brain cells of the *post-mortem* human brain, as characterized across five different studies. For PLS1+, we found significant enrichment in genes typically expressed in microglia, astrocytes and oligodendrocytes precursor cells (Figure 7B and Supplementary Table S4). For PLS1-, we found significant enrichment in genes typically expressed in excitatory and inhibitory neurons (Figure 7B and Supplementary Table S2).

### Enrichment for genes related to pain and other brain disorders

Finally, we investigated whether PLS1+ and PLS1- are enriched for pain- and other brain disorders-related genes as identified in previous studies. For PLS1+, we found enrichment for pain-related (OR = 1.40, p = 0.04), but not genes related to any of the other brain disorders we tested (Supplementary Table S3). For PLS1-, we did not find enrichment for pain-related genes (OR = 1.07, p = 0.42), but found enrichment for genes associated with epilepsy (OR = 1.54, p = 0.02, p_FDR_ = 0.09) and major depressive disorder (OR = 1.57, p = 0.02, p_FDR_ = 0.09) (Supplementary Table S3).

## Discussion

In this manuscript, we investigated changes in cortical MS in three independent cohorts of patients with chronic pain syndromes, i.e. knee osteoarthritis, chronic low back pain and fibromyalgia. Moreover, we leveraged transcriptomic data of brain-wide gene expression in the *post-mortem* human brain to uncover patterns of gene expression that are aligned with the changes in cortical MS observed across these pain conditions. Our study yielded two key findings. First, we uncovered a new and robust pattern of cortical MS remodelling across three chronic pain syndromes, which was different from that observed in patients with MDD and points towards the existence of shared disease-mechanisms driving cortical remodelling that cut across the boundaries of specific pain syndromes. Second, we demonstrate that cortical MS remodelling in chronic pain spatially correlates with the brain-wide expression of genes involved in glial immune response and neuronal plasticity, which we suggest might drive divergent tails of MS cortical remodelling during chronic pain. These findings might explain the partial efficacy of existing therapies and provide food for thought in how future treatment development might be pursued. We discuss each of these main findings below in detail.

Morphometric similarity mapping disclosed a robust pattern of cortical MS changes across chronic pain syndromes, which generally involved small-to-medium sized increases in the insula and limbic cortices, and decreases in the occipital, sensorimotor and frontal cortices. These findings are interesting for several reasons. First, of the various brain regions that have been implicated in the perception of pain, the insula and limbic system are among the ones most consistently reported across studies^45^. Second, functional and structural alterations in these regions have often been reported in neuroimaging studies of patients with different chronic pain syndromes^9,17-19^, including increases in the connectivity of the insula with nodes of the default-mode network across multiple chronic pain syndromes^46-48^. Third, previous studies have further demonstrated that the limbic system plays a key role in the transition from acute to chronic pain^49,50^. Altogether, these aspects reinforce the neuroanatomical plausibility of the pattern in regional MS changes we report in this manuscript.

What does this MS phenotype represent? Morphometric similarity quantifies the correspondence or kinship of two cortical areas in terms of multiple macrostructural features (e.g., cortical thickness) that are measurable by MRI (note we only included structural data, but MS can easily accommodate data from other modalities such as microstructure features)^33^. Hence, high MS between a pair of cortical regions indicates that there is a high degree of correspondence between them in terms of cytoarchitectonic features that we cannot directly observe given the limited spatial resolution and lack of cellular specificity of MRI. This assumption has received empirical support in prior work showing that morphometrically similar cortical regions share patterns of gene co-expression and are more likely to be axonally connected to each other^33^. Therefore, here we interpret the reduced MS that we observed over frontal, occipital and sensorimotor brain regions in chronic pain as indicating that there is reduced cytoarchitectonic similarity, or greater cytoarchitectonic differentiation, between these areas and the rest of the cortex, which is probably indicative of reduced anatomical connectivity to and from the less similar, more differentiated cortical areas. On the other hand, increased morphometric similarity in the insula and limbic regions implies increased cytoarchitectonic similarity and, perhaps, axonal connectivity with the rest of the cortex.

Interestingly, we found a significant negative association between MS changes in patients and MS in healthy controls: regions with typically low MS in healthy controls tend to increase MS in chronic pain patients, while regions with high MS in healthy controls tend to decrease MS in patients. This shows that the typical morphometric organization of a cortical region might determine the output of cortical remodelling during chronic pain. Following from the interpretation above, this association points to two key patterns of changes in brain MS during chronic pain: decoupling of hub regions and dedifferentiation of highly differentiated regions. Previous fMRI and DTI brain networks studies have demonstrated that “hub” regions are more likely to be disturbed and reduce their connectivity in the presence of brain disease^44^. Therefore, it is tempting to speculate that the decreases in MS during chronic pain we describe here might reflect an overall pattern of decreases in axonal connectivity of “hub” regions with the rest of the cortex, as observed across other brain disorders. The reverse, i.e. increased connectivity, might drive increases in MS during chronic pain. Nevertheless, we cannot exclude that either increases or decreases in MS might simply reflect local changes in cytoarchitectonics or even a combination of local tissue changes and connectivity. Whether these changes are permanent or might be reversible after treatment is a question that should be investigated in future longitudinal studies. Previous studies have reported structural changes in the brain of patients with chronic pain that reverted after treatment^51-53^. Hence, it is not implausible that the changes in MS we report here might attenuate or even revert once effective pain control is achieved.

In an attempt of connecting these MS changes during chronic pain to the gene expression and cellular pathways potentially driving those changes, we used PLS to identify the weighted combination of genes in the whole transcriptome that has a cortical expression map most similar to the cortical map of cross-condition case–control MS differences we derived for chronic pain patients. We found 338 genes (PLS1+) that are highly expressed in regions where MS increases and 236 genes (PLS1-) highly expressed in regions where MS decreases. The proteins encoded by these genes are part of topologically clustered interaction networks that were significantly enriched for a number of relevant GO biological processes and KEGG pathways. In PLS1+, we found predominance of genes related to the glial immune response and highly expressed in microglia, astrocytes and oligodendrocyte precursor cells. Further reinforcing the relationship of this subset of genes with pain, we found enrichment in PLS1+ for pain-related genes but not genes related other brain disorders. In PLS1-, we found predominance of genes related to calcium signalling and long-term potentiation, that are highly expressed in excitatory and inhibitory neurons. Interestingly, here, we did not find enrichment for pain-related genes, but found enrichment for genes related to epilepsy and major depressive disorder. This last finding is in keep with the idea that chronic pain might share neurobiological pathways with epilepsy ^54-56^ and depression^42^. Furthermore, it matches well the clinical observation that antiepileptic drugs^57^ and antidepressants^58^ (drugs used to treat epilepsy and depression, respectively) also improve pain in patients with chronic pain syndromes. Altogether, these findings suggest that alterations in glial immune response and neuronal plasticity, both key elements of the current pathophysiological models of chronic pain^59,60^, might drive divergent aspects of MS cortical remodelling during chronic pain.

Mounting preclinical evidence has demonstrated that neuroinflammation, characterized by infiltration of immune cells, activation of resident glial cells and production of inflammatory mediators in the peripheral and central nervous system, induces pain and contributes to its persistence during chronic pain^61^. The overproduction of proinflammatory cytokines and chemokines, and other mediators, are thought to be responsible for neuronal sensitization of pain pathways, at both spinal and brain levels, with subsequent maintaining of pain^59^. In humans, the neuroinflammation hypothesis has received direct support from positron emission tomography (PET) studies showing increased binding of ligands for the 18-kDa translocator protein (TSPO), currently used as a marker of neuroinflammation and glial activation in the brain^62-66^, spinal cord and nerve roots^64^ of patients with chronic pain as compared to healthy controls. Our transcriptomic – imaging association findings for PLS1+ are broadly compatible with this hypothesis. In chronic pain, MS increases predominantly in regions of low MS in healthy controls. As explained above, this implies that highly differentiated regions of the cortex undergo morphometric dedifferentiation during chronic pain. Based on our transcriptomic association findings, we suggest that neuroinflammation, with neuronal loss, synapse removal and glial proliferation^67^, might disrupt cortical organization and drive loss of cortex differentiation during chronic pain^68,69^.

Maladaptive neuronal plasticity, with dysfunctional regulation of the cortical E/I balance has similarly been suggested to contribute to chronic pain^69-72^ and is targeted by at least some of the drugs currently used to treat chronic pain patients, such as anticonvulsants (i.e. gabapentinoids)^73^. Indeed, shifting of the E/I towards hyperexcitability, as a consequence of either enhanced excitation or reduced inhibition, is thought to augment central pain processing^69,72^. One key idea around this model is the hypothesis that peripheral injury triggers plastic changes or long-term potentiation (LTP) in the cortical synapses. Such potentiation or excitation persists for a long period of time, and consequently might generate abnormal neuronal spike activity in the brain without obvious peripheral sensory stimulation^60^. This model speaks directly to our transcriptomic – imaging association findings for PLS1-, where we found enrichment in genes related to LTP and calcium signalling, and highly expressed in excitatory and inhibitory neurons. How could alterations in E/I and LTP explain morphometric differentiation (decreases in MS) of cortical regions which typically exhibit low differentiation (high MS)? LTP promotes the formation of synapses and remodelling of dendritic spine substructures^74^. Dendritic spines are morphological specializations that receive synaptic inputs and compartmentalize calcium^74^. Based on our transcriptomic association findings, we suggest that changes in synaptic plasticity of excitatory and inhibitory neuronal populations regulating E/I could drive modifications in cortical microstructure and reorganization, which ultimately at the long-term might manifest macroscopically as an increase in MS^69^. However, as for our model above on neuroinflammation-related MS increases, we should acknowledge that, for now, these hypotheses remain speculative. Future studies combining ex-vivo MRI and histological examinations of the post-mortem human brain of patients with chronic pain might help testing our hypotheses further.

Our study has some limitations. First, the whole-brain gene expression data derives only from six *post-mortem* adult brains (mean age = 43 y) and not from patients with chronic pain. Furthermore, the AHBA only includes data in the right hemisphere from two donors, which led us to exclude MS-changes in the right hemisphere for the transcriptomic association analyses. Second, we pooled data from three different cohorts of patients with different chronic pain syndromes that were collected using different protocols and setups. This aspect poses limitations for investigating condition-specific changes in MS, which are likely to exist and might be interesting to pursue. Indeed, even though in the PCA we found a dominant component explaining around 65%, this left 35% of unexplained variance which might pertain to patterns of MS changes characteristic of each of these conditions. These condition-specific changes should ideally be investigated in a larger study assessing all patients under the same protocol to avoid study-specific confounds. Moreover, even within the boundaries of a specific chronic pain syndrome, it is likely that different pathophysiological mechanisms are in play in different patients^75-77^. This within-group heterogeneity is an aspect we did not deal with in this study, but that future studies should take into consideration. We note though that by including three distinct chronic pain conditions which are typically treated with different pharmacological/interventional strategies, we reduce the potential confound of past/current treatment. Fourth, and following from the latter, the datasets have varied, limited clinical information available, making it difficult to assess the clinical significance of the morphometric similarity phenotype. Fifth, the MDD cohort we used to define the pattern of MS changes was smaller (N=19) and not age-matched to our chronic pain cohorts. Even though in all of our MS analyses we regressed out the effects of age, we cannot exclude that MDD in older ages might manifest with a different MS phenotype, which might be more similar to that we report here for chronic pain. This seems unlikely though, given that our MDD and CLBP cohorts had very similar mean ages and still we found a negative correlation between MS changes in these conditions. Finally, while our findings are suggestive of a contribution of neuroimmune responses and neural plasticity to the changes in MS we report here, our correlational approach does not allow us to infer a causal role of these pathways. Future longitudinal studies might shed further light on the link between these transcriptomic pathways and progressive changes in MS in chronic pain patients by investigating whether pharmacologic modulation of either pathway might attenuate the changes in MS we report here.

In summary, our study describes a new and robust pattern of cortical MS remodelling across three chronic pain syndromes and identifies alterations in the glial immune response and imbalances in neuronal plasticity as candidates for molecular and cellular mechanisms driving divergent tails of these cortical changes. Altogether, our data indicates that chronic pain entails a shared component of disease-mechanisms that goes beyond specific clinical syndromes boundaries and might involve disruption of multiple elements of the cellular architecture of the brain which is unlikely to be efficiently targeted by current one-size-fits-all treatments. Ultimately, these findings highlight that developing new effective therapeutic approaches to the brain pathology that accompanies chronic pain might require a multi-target approach modulating both glial function and neuronal plasticity.

## Methods

### Samples

We used structural MRI data from three prior case–control studies on knee osteoarthritis (OA)^78^, chronic low back pain (CLBP)^49^ and fibromyalgia^79^. The full details of each sample (including inclusion and exclusion criteria) have been described in the original publications. The OA dataset included 56 OA chronic pain patients (54% women, 58 ± 6.96 y) and 20 age-matched healthy control subjects (50% women, 58 ± 6.65 y). The CLBP dataset included 29 CLBP patients (59% women, 30.79 ± 11.59 y) and 33 healthy controls (14 females, 31.18 ± 9.65 y). The fibromyalgia dataset included 20 women with fibromyalgia (46.4 ± 12.4 y) and 20 age-matched healthy control women (42.1±12.5 y). The original studies were approved by the competent ethics assessment board of each respective host institution. All participants provided informed consent before enrolling the respective studies.

### MRI data acquisition

We used high-resolution T1-weighted anatomical images acquired in 3T scanners. We provide a summary of the parameters used for T1-weighted anatomical images acquisition in each dataset (as described in the original publications).

### OA dataset

images were acquired in a Siemens 3.0 Trio whole-body scanner using the standard radio-frequency head coil with the following parameters: TR/TE = 2500/3.36 ms; flip angle = 9°; in-plane matrix resolution = 256 × 256; FOV = 256 mm^2^; slice thickness = 1 mm, number of slices = 160^78^.

### CLBP dataset

images were acquired in a Siemens 3.0 T Trio B whole-body scanner equipped with a 32-channel head coil with the following parameters: TR/TE = 1900/2.52 ms, flip angle = 9°, matrix 256 × 256, slice thickness = 1 mm, number of slices = 176^49^.

### Fibromyalgia dataset

images were acquired in a 3.0 Tesla GE Discovery MR750 scanner (HD, General Electric Healthcare, Waukesha, WI, USA) and a commercial 32-channel head coil array, using the FSPGR BRAVO pulse sequence: TR/TE = 7.7/3.2 ms, flip angle = 12°, matrix = 256 × 256, FOV = 256 mm^2^, slice thickness = 1 mm, number of slices = 168^79^.

### Morphometric Similarity (MS) Mapping

The T1-weighted MRI data from all participants were preprocessed using the recon-all command from FreeSurfer (version 6.0)^80^. The surfaces were then parcellated using the 68 cortical regions of the Desikan–Killiany (DK) atlas^**81**^. For each cortical region, we estimated five parameters: gray matter volume, surface area, cortical thickness, gaussian curvature and mean curvature. Each parameter was normalized for sample mean and standard deviation to account for variation in value distributions between the features. After normalization, MS networks were generated by computing the regional pairwise Pearson correlations in morphometric feature sets, yielding an a 68×68 morphometric similarity matrix *ℳ*_*i*_ for each participant, *i*=1, …, *N*, which represents the strength of MS between each pair of cortical areas. For all individuals, regional MS (i.e., nodal similarity) estimates were calculated as the average MS between a given cortical region and all others. While MS calculation allows for the inclusion of data from different imaging modalities, here we used only features extracted from T1-weighted MRI data. It has been previously demonstrated that there is high spatial concordance (*r* = 0.91) between the topography of regional MS derived from T1-weighted MRI data alone, and regional MS from more modalities (e.g. a combination of T1- and diffusion weighted imaging)^33^.

### Case–Control Analysis of Morphometric Similarity Networks

The global MS for each participant is the average of *ℳ*_*i*_. The regional *MS*_*i,j*_ for the i^th^ participant at each region *j*=1,…,68 is the average of the j^th^ row (or column) of *ℳ*_*i*_. Thus, regional MS is equivalent to the weighted degree or “hubness” of each regional node, connected by signed and weighted edges of pair-wise similarity to all other nodes in the whole brain connectome represented by the MS matrix. For global and regional MS statistics alike, we fit linear models to estimate case–control differences, with age, sex, and total intracranial volume as covariates. The resulting *P* value for each region was thresholded for significance using *FDR*<0.05, to control type 1 error over multiple (n = 68) tests. To test for the differential contribution of single cortical features to the observed regional MS changes in each chronic pain condition, we recomputed condition-specific MS change maps with exclusion of each individual cortical feature prior to MS calculation and then determined which of these single-feature exclusions most change the topography of the observed MS changes. We did so by calculating pairwise Spearman correlations between the original and all other leave-one-out maps of MS changes, in each condition separately.

### Cortical morphometric similarity remodelling in Major Depressive Disorder and its association with chronic pain

Chronic pain is often comorbid with major depressive disorder (MDD)^**41**^ and these two entities share brain mechanisms of neuroplasticity^**42**^. We used another openly available dataset (see the original article for further details^43^) including high resolution structural brain data from 19 unmedicated patients with MDD (11 females, 29.45 ± 11.26 y) and 20 age-matched healthy controls (11 females, 33.52 ± 14.07 y). We characterized changes in regional MS associated with MDD and investigated whether changes in regional MS in MDD can predict those we observed in chronic pain patients. We calculated MS using the procedures described above and tested for case-control differences between MDD patients and healthy controls using linear models, where we accounted for age, sex and intracranial volume. Finally, we calculated pairwise Spearman correlations between changes in regional MS in MDD and those observed in the different chronic pain conditions.

### Mapping case-control differences in regional morphometric similarity to established patterns of cytoarchitectonic cortical organization

To help us to contextualize the case-control differences in regional MS we observed for the different chronic pain conditions, we mapped them in relation to well established patterns of cytoarchitectonic organization of the cortex. To that end, we used the *von Economo* atlas of cortex classified by cytoarchitectonic criteria^37^. For each subject, we quantified MS within each parcel of these atlas and then performed case-control comparisons using linear models, with age, gender and intracranial volume as covariates.

### Morphometric similarity hubs susceptibility

We investigated relationships between case-control differences in regional MS and the typical pattern of regional MS distribution in healthy controls with Spearman correlations. In keeping with histological results indicating that cytoarchitectonically similar areas of the cortex are more likely to be anatomically connected and that morphometric similarity in the macaque cortex was correlated with tract-tracing measurements of axonal connectivity^**33**^, we followed the approach suggested by *Seidlizt et al*.^32^ to map each region to one of four patterns of changes in MS: (i) regions of low MS in healthy controls (highly differentiated from the rest of the cortex) that increase their MS during with the rest of the cortex during chronic pain (De-differentiation); (ii) regions of high MS in healthy controls (highly connected with the rest of the cortex) that increase their MS during chronic pain (Hypercoupling); (iii) regions of low MS in healthy controls that decrease their MS during chronic pain (Hyper-differentiation); and (iv) regions of high MS in healthy controls that decrease their MS during chronic pain (Decoupling). We subdivided each axis of the scatter plot in two sections, one above and another below 0, which resulted in four quadrants, each representing one of the four scenarios presented above. We then quantified the percentage of regions falling within each of these four quadrants to identify dominant patterns of change.

### Defining a cross-condition pattern of changes in regional morphometric similarity during chronic pain

We investigated the similarity in case-control changes in regional MS across chronic pain conditions by calculating pairwise Spearman correlations of regional case-control statistics (Z-scores) from each condition^39^. We found significant correlations for all possible pairs of conditions, which indicated that remodelling of regional MS during chronic pain shares a common pattern across conditions. We then ran principal component analysis (PCA) on the three vectors of case-control changes in regional MS to identify this shared profile of cross-condition changes. The first component alone explained 64.45% of the shared variance, the second 25.38% and the third 10.16%. Only the first component showed an eigenvalue > 1 (1.93). Hence, we kept only the first PC since case-control changes in regional MS across our three chronic pain conditions seem to be well captured by one single dominant cross-condition pattern.

### Microarray expression data: Allen Human Brain Atlas (AHBA)

Regional microarray expression data were obtained from six post-mortem brains provided by the Allen Human Brain Atlas (AHBA; http://human.brain-map.org/) (ages 24–57 years)^82^. We used the *abagen* toolbox (https://github.com/netneurolab/abagen) to process and map the transcriptomic data to 84 parcellated brain regions from the DK atlas^81^. Briefly, genetic probes were reannotated using information provided by Arnatkeviciute et al., 2019^83^ instead of the default probe information from the AHBA dataset, hence discarding probes that cannot be reliably matched to genes. Following previously published guidelines for probe-to-gene mappings and intensity-based filtering^83^, the reannotated probes were filtered based on their intensity relative to background noise level; probes with intensity less than background in ≥50% of samples were discarded. A single probe with the highest differential stability, ΔS(p), was selected to represent each gene, where differential stability was calculated as^84^:

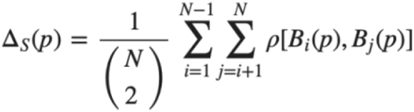

Here, ρ is Spearman’s rank correlation of the expression of a single probe *p* across regions in two donor brains, Bi and Bj, and *N* is the total number of donor brains. This procedure retained 15,633 probes, each representing a unique gene.

Next, tissue samples were assigned to brain regions using their corrected MNI coordinates (https://github.com/chrisfilo/alleninf) by finding the nearest region within a radius of 2 mm. To reduce the potential for misassignment, sample-to-region matching was constrained by hemisphere and cortical/subcortical divisions. If a brain region was not assigned to any sample based on the above procedure, the sample closest to the centroid of that region was selected in order to ensure that all brain regions were assigned a value. Samples assigned to the same brain region were averaged separately for each donor. Gene expression values were then normalized separately for each donor across regions using a robust sigmoid function and rescaled to the unit interval. Scaled expression profiles were finally averaged across donors, resulting in a single matrix with rows corresponding to brain regions and columns corresponding to the retained 15,633 genes. As a further sanity check, we conducted leave-one-donor out sensitivity analyses to generate six expression maps containing gene expression data from all donors, one at a time. The principal components of these six expression maps were highly correlated (average Pearson correlation of 0.993), supporting the idea that our final gene expression maps where we averaged gene expression for each region across the six donors is unlikely to be biased by data from a specific donor. Since the AHBA only includes data for the right hemisphere for two subjects, in our transcriptomic-imaging association analyses we only considered the left hemisphere cortical regions (34 regions).

### Identifying transcriptomic correlates of cortical morphometric similarity remodelling in chronic pain

In order to be able to investigate associations between cross-condition changes in MS during chronic pain and brain gene expression, we used partial least square regression (PLS)^31^. Partial least square regression uses the gene expression measurements (the predictor variables) to predict the regional MS changes (the response variables). This approach allows us to rank all genes by their multivariate spatial alignment with cross-condition regional MS changes during chronic pain. The first PLS component (PLS1) is the linear combination of the weighted gene expression scores that have a brain expression map that covaries the most with the map of MS changes. As the components are calculated to explain the maximum covariance between the dependent and independent variables, the first component does not necessarily need to explain the maximum variance in the dependent variable. However, as the number of components calculated increases, they progressively tend to explain less variance in the dependent variable. Here, we tested across a range of components (between 1 and 15) and quantified the relative variance explained by each component. The statistical significance of the variance explained by each component was tested by permuting the response variables 1,000 times.

In our analysis, a solution with a single component explained variance in regional MS changes above chance (p_boot_=0.003). The first PLS component (PLS1) alone explained the highest amount of variance alone (24.42%). Hence, we focused our further gene set enrichment analyses on PLS1. The error in estimating each gene’s PLS1 weight was assessed by bootstrapping (resampling with replacement of the 34 brain regions), and the ratio of the weight of each gene to its bootstrap standard error was used to calculate the *Z* scores and, hence, rank the genes according to their contribution to PLS1^37^. Genes with large positive PLS1 weights correspond to genes that have higher than average expression in regions where MS increases, and lower than average expression in regions where MS decreases. Mid-rank PLS weights showed expression gradients that are weakly related to the pattern of MS changes. On the other side, genes with large negative PLS1 weights correspond to genes that have higher than average expression in regions where MS decreases the most, and lower than average expression in regions where MS increases. Hence, from the ranked PLS1 list of genes, we then selected all genes with positive and negative weights Z>3 and Z < -3, respectively (all p_FDR_< 0.05, FDR corrected for the total number of genes tested). PLS1 genes with Z>3 are for simplicity termed PLS1+ and genes with Z<-3 PLS1-. While our choice of Z = 3 as a threshold to identify the most positively and negatively genes associated with MS changes in chronic pain patients is somehow arbitrary, we note that Z=3 in our case corresponds to a stringent threshold of p_FDR_ = 0.036 (more stringent than p_FDR_ < 0.05). We used these two sets of genes for further enrichment analyses, as described below. We confirmed our enrichment analyses were not driven by the choice of this specific threshold by repeating all analyses described below considering all genes that passed a more liberal threshold of p_FDR_<0.05. The overall pattern of results did not change.

### Protein-Protein networks and gene set enrichment analysis

We then used all genes in PLS1+ and PLS1-to conduct further bioinformatics analyses investigating whether these genes map to common and relevant biological pathways. First, we used *STRING* (version 10.5)^85^ to construct protein-protein functional interactions networks. We excluded text mining as an active interaction source and used the default medium required interaction score of 0.400 to identify all possible links within our list of target genes. Second, we used the *GENE2FUNC* function from the Functional Mapping and Annotation of Genome-Wide Association Studies (*FUMA*)^86^ platform to investigate functional enrichments using rank-based gene ontology (GO) and KEEG pathways enrichment analysis. We used as background all AHBA genes that passed our pre-processing criteria and hence were used in our PLS analyses (n=15,633). We corrected for multiple gene-set enrichment testing by applying FDR correction.

### Brain cell-type enrichment analysis

We also investigated whether our PLS1+ and PLS1-subsets of genes was particularly enriched for genes of specific brain cell types. We compiled data from five different single-cell studies using *post-mortem* cortical samples in human postnatal subjects to avoid any bias based on acquisition methodology or analysis or thresholding^87-91^. To obtain gene sets for each cell type, categorical determinations were based on each individual study, as per the respective methods and analysis choices in the original paper. All cell-type gene sets were available as part of the respective papers. We generated a single omnibus gene list for each cell type by merging the study-specific gene lists, and then filtered it to retain only genes sampled in the AHBA. Two studies did not subset neurons into excitatory and inhibitory^87,90^, and thus those gene sets were excluded from the cell-class assignment. Additionally, only one study included the annotation of the “Per” (pericyte) type, and thus we did not consider that cell type^88^. This approach has already been validated elsewhere^32^.

We then conducted cell-class enrichment analyses using the *GeneOverlap* package from R (version 1.26.0). We used as background all AHBA genes that passed our pre-processing criteria and hence were used in our PLS analysis (n=15,633). *GeneOverlap* calculates the overlap between two sets of genes (in our case, the set of PLS1+ or PLS1-, and each of the brain cell type omnibus gene sets derived as explained above) and uses a Fisher’s exact test to find whether the overlap between these two sets is higher than one would expect by randomly selecting a subset of genes from the background with the same number of elements. Here, enrichment is quantified as an odds ratio, where values lower or equal to 1 indicate minimal overlap between sets and hence absence of enrichment. Therefore, the null hypothesis is that the odds ratio is no larger than 1. Significant odds ratio larger than 1 indicate enrichment for genes of a specific cell type. We applied FDR correction for the number of cell types tested.

### Pain- and other brain disorders-related genes enrichment

We also investigated whether our PLS1+ and PLS1-subsets of genes were particularly enriched for pain- and other brain disorders-related genes. The list of previously identified pain-related genes was defined using the following public resources: 1) the pain-related genes identified in mice gene knockout studies collected in the Pain Gene Database (http://paingeneticslab.ca/4105/06_02_pain_genetics_database.asp)^92^, 2) the pain-related genes identified in humans collected in the Human Pain Genes Database (https://humanpaingenetics.org/hpgdb)^93^, 3) the Pain Research Forum (https://www.painresearchforum.org/resources/pain-gene-resource)^94^, 4) the genes involved in human pain diseases collected in the DisGeNET (http://www.disgenet.org)^95^, and 5) the genes considered to be functioning in pain perception summarized in Gene Ontology (GO:0019233). In total, we identified 2111 pain-related genes included in the above public resources. 807 of these genes were not part of our initial list of 15633 AHBA genes that passed our pre-processing criteria and were excluded from further analyses (please see Supplementary data S3 for the list of 1304 pain-related genes used in these analyses). The lists of genes associated with other brain disorders (alzheimer’s disease, parkinson disease, huntington’s disease, epilepsy, autism spectrum disorder, major depressive disorder, anxiety, bipolar disorder and schizophrenia) were collected from the DisGeNET. We tested for gene enrichment in both PLS1+ and PLS1-subsets using *GeneOverlap*, as explained above.

### Spatial permutation test (spin test)

In several analyses in the current study, we investigated the spatial correspondence between different imaging-derived measures. While several studies have reported significance based on the assumption that the number of samples is equal to the number of regions, this is technically inaccurate, as the number of regions is both arbitrary (due to the resolution of the chosen parcellation) and non-independent (due to spatial autocorrelation amongst neighbouring parcels). To overcome this issue, we used spatial permutation tests (spin test) as implemented in previous studies^29,96,97^. This approach consists in comparing the empirical correlation amongst two spatial maps to a set of null correlations, generated by randomly rotating the spherical projection of one of the two spatial maps before projecting it back on the brain surface. Importantly, the rotated projection preserves spatial contiguity of the empirical maps, as well as hemispheric symmetry. Therefore, each analysis correlating values from two cortical maps is reported with a *P*-value derived from the spherical permutation (*P*_spin_), obtained by comparing the empirical Spearman’s Rho to a null distribution of 10 000 Spearman correlations, between one empirical map and the randomly rotated projections of the other map. The matlab code to implement this test can be found in https://github.com/frantisekvasa/rotate_parcellation.

## Data availability

Data can be accessed from open repositories in the following links (Osteoarthritis: https://openneuro.org/datasets/ds000208/versions/1.0.0; Chronic Low Back Pain: http://www.openpain.org; Fibromyalgia: https://openneuro.org/datasets/ds001928/versions/1.1.0; Major Depressive Disorder: https://openneuro.org/datasets/ds000171/versions/00001). A reporting summary for this Article is available as a Supplementary Information file.

## Code availability

The code for morphometric similarity and gene expression association analyses is available at https://github.com/SarahMorgan/Morphometric_Similarity_SZ.

## List of Supplementary Materials

Supplementary Figure S1. Regional distribution of morphometric similarity in healthy controls.

Supplementary Figure S2. Correlations between the distributions of regional morphometric similarity in healthy controls across datasets.

Supplementary Figure S3. Correlations between case-control differences in regional morphometric similarity across chronic pain conditions.

Supplementary Figure S4. Leave-one-feature-out analysis.

Supplementary Figure S5. Leave-one-feature-out analysis (cortical maps)

Supplementary Figure S6. Cortical morphometric similarity remodelling in major depressive disorder and its association with cortical morphometric similarity remodelling in chronic pain

Supplementary Table S1. Chronic pain-related changes in regional morphometric similarity within the parcels of the *Von Economo* atlas of the cortex

Supplementary Figure S7. STRING protein-protein networks

Supplementary Table S4. Cell-type enrichment analyses (full statistics)

Supplementary Data S1. Full ranked list of PLS1 genes

Supplementary Data S2. FUMA gene-set enrichment analysis (full results)

Supplementary Data S3. List of pain-related genes

## Acknowledgments

We would like to thank all volunteers contributing data to this study and all researchers making their data openly available.

## Author contributions

DM designed the study, performed the data analysis and drafted the manuscript; all authors discussed the findings, revised the manuscript for intellectual content and approved the final version of the manuscript.

