## Supplementary for "Transcriptional and cellular signatures of cortical morphometric similarity remodelling in chronic pain"

Martins et al.

**Supplementary Figure S1. Regional distribution of morphometric similarity in healthy controls.** In this figure, we provide cortical maps depicting the distribution of morphometric similarity values in the cortex of healthy controls from each dataset.

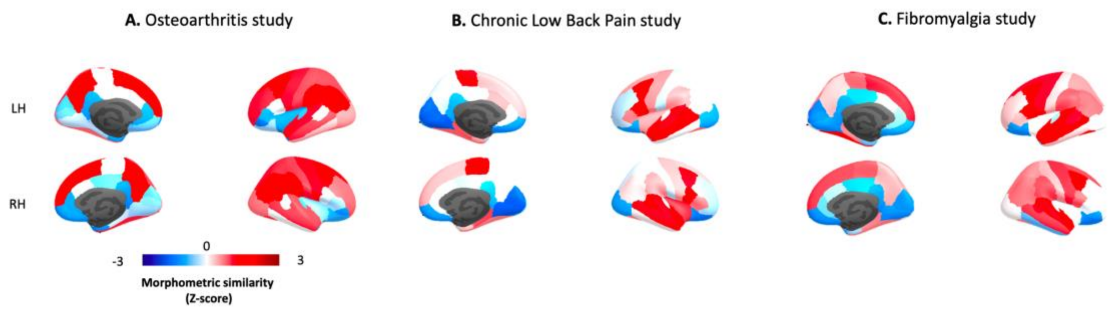


**Supplementary Figure S2. Correlations between the distributions of regional morphometric similarity in healthy controls across datasets.** In this heatmap, we show pairwise spearman correlations between normalized (Z-scores) values of regional morphometric similarity in the cortex of healthy controls from each dataset. Significance was assessed with spatial permutation testing. $^{***}p_{spin} < 0.001$. *Abbreviations*: OA – Osteoarthritis; CLBP – Chronic low back pain.

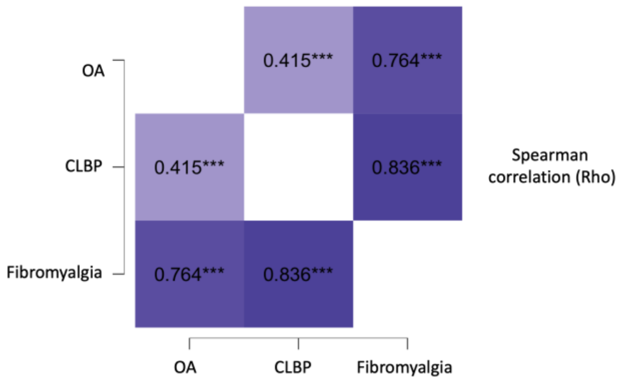


**Supplementary Figure S3. Correlations between case-control differences in regional morphometric similarity across chronic pain conditions.** In this heatmap, we show pairwise spearman correlations between normalized (Z-scores) values of regional morphometric similarity changes in each chronic pain condition. Significance was assessed with spatial permutation testing. $*p_{spin} < 0.05$; $^{***}p_{spin} < 0.001$.

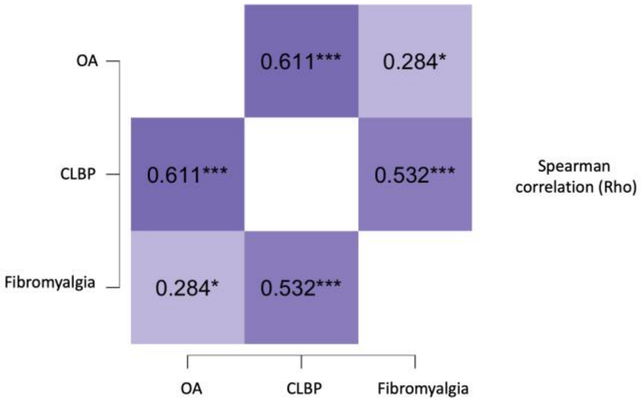


**Supplementary Figure S4. Leave-one-feature-out analysis.** In these heatmaps, we show the results of our leave-one-feature-out analyses testing for differential contribution of single cortical features to the observed regional morphometric similarity (MS) changes in each chronic pain condition. Briefly, we recomputed condition-specific MS change maps excluding each individual cortical feature at a time prior to MS calculation. We then pairwise spearman correlations between normalized (Z-scores) values of regional MS changes of each leave-one-feature out, including our map calculated using all features (all). Significance was assessed with spatial permutation testing. $^{**}p_{spin} < 0.01$; $^{***}p_{spin} < 0.001$. *Abbreviations*: GC – Gaussian curvature; GMV – Grey matter volume; MC – Mean curvature; CT – Cortical thickness.

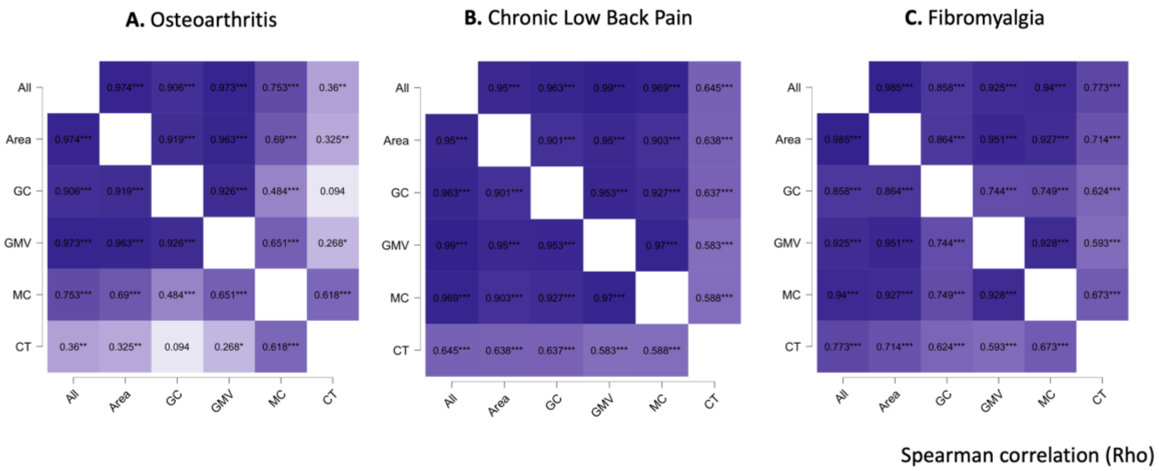


**Supplementary Figure S5. Leave-one-feature-out analysis (cortical maps).** In these cortical maps, we show recomputed condition-specific MS change maps excluding each individual cortical feature at a time prior to MS calculation. *Abbreviations*: OA – Osteoarthritis; CLBP – Chronic low back pain; FM – Fibromyalgia; GC – Gaussian curvature; GMV – Grey matter volume; MC – Mean curvature; CT – Cortical thickness.

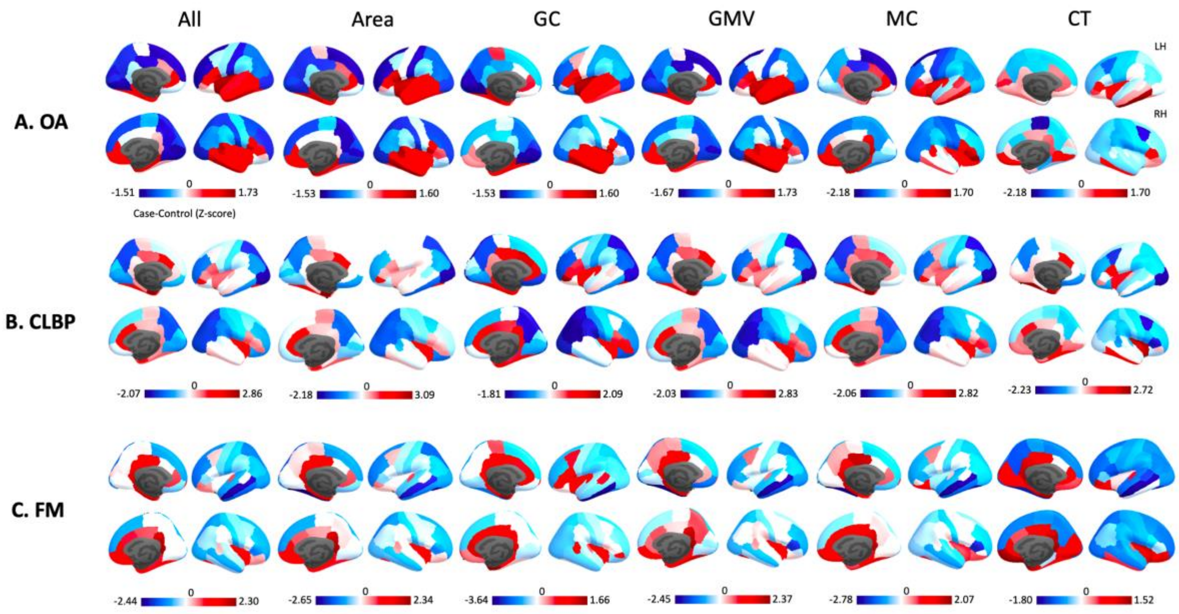


**Supplementary Figure S6. Cortical morphometric similarity remodelling in major depressive disorder and its association with cortical morphometric similarity remodelling in chronic pain.** In panel A, present a cortical map depicting changes in regional morphometric similarity (MS) in major depressive disorder (MDD). In panel B, we present a heatmap showing the pairwise spearman correlations between changes in regional MS in MDD and those observed in the different chronic pain conditions. Note that in addition to testing for correlations with regional MS changes in each chronic pain condition, we also included PCA1 which represent cross-condition changes as captured in a principal component analysis run across the three conditions. Significance was assessed with spatial permutation testing. $^{*}p_{spin} < 0.05$; $^{**}p_{spin} < 0.01$; $^{***}p_{spin} < 0.001$. *Abbreviations*: OA – Osteoarthritis; CLBP – Chronic low back pain; FM – Fibromyalgia.

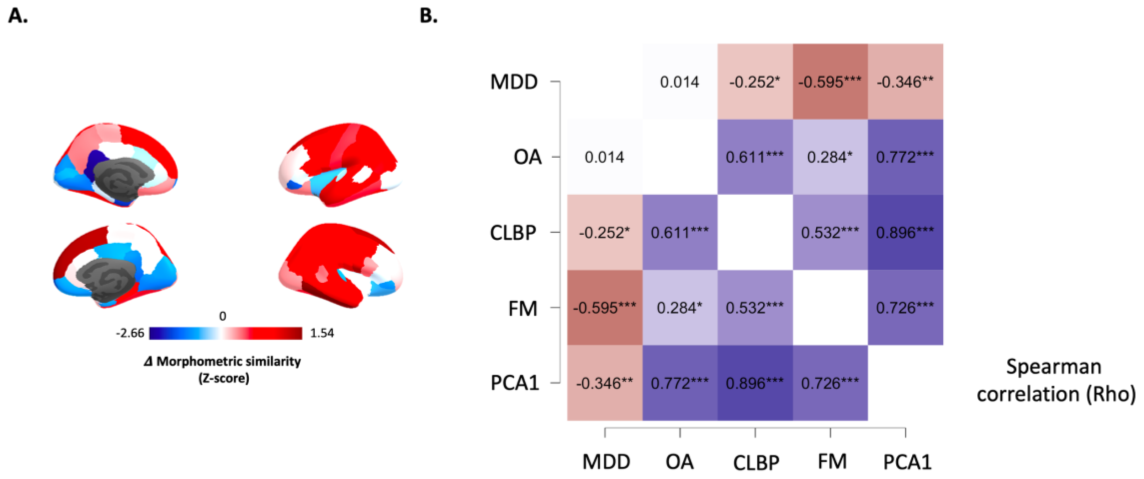

**Supplementary Table S1. Chronic pain-related changes in regional morphometric similarity within the parcels of the *Von Economo* atlas of the cortex.** This table summarizes the full statistics of the analyses run on each parcel of the *Von Economo* atlas of the cortex classified according to cytoarchitectonic criteria. *Abbreviations:* OA – Osteoarthritis; CLBP – Chronic low back pain.

| Condition | Statistics | Agranular, primary motor | Association A | Association B | Secondary sensory | Primary Sensory | Limbic | Insular |
|---|---|---|---|---|---|---|---|---|
| **OA** | **T-stats** | -0.551 | 1.052 | 0.051 | -1.158 | -1.955 | 1.406 | 2.376 |
| | **Cohen's d** | -0.13 | 0.25 | 0.01 | -0.27 | -0.46 | 0.33 | 0.56 |
| | **$P_{uncorrected}$** | 0.58 | 0.30 | 0.96 | 0.25 | 0.06 | 0.16 | 0.02 |
| | **$P_{FDR}$** | 0.68 | 0.42 | 0.96 | 0.42 | 0.21 | 0.37 | 0.14 |
| **CLBP** | **T-stats** | 0.081 | 0.091 | -1.240 | -0.999 | -1.279 | 2.172 | 1.655 |
| | **Cohen's d** | 0.02 | 0.02 | -0.33 | -0.26 | -0.34 | 0.57 | 0.44 |
| | **$P_{uncorrected}$** | 0.94 | 0.93 | 0.22 | 0.32 | 0.21 | 0.04 | 0.10 |
| | **$P_{FDR}$** | 0.94 | 0.94 | 0.39 | 0.49 | 0.39 | 0.28 | 0.35 |
| **Fibromyalgia** | **T-stats** | -0.43 | -2.30 | -1.01 | 0.09 | -0.70 | 2.72 | 0.77 |
| | **Cohen's d** | -0.14 | -0.77 | -0.34 | 0.03 | -0.23 | 0.91 | 0.26 |
| | **$P_{uncorrected}$** | 0.67 | 0.03 | 0.32 | 0.92 | 0.49 | 0.01 | 0.45 |
| | **$P_{FDR}$** | 0.78 | 0.11 | 0.69 | 0.92 | 0.69 | 0.07 | 0.69 |

**Supplementary Figure S7. STRING protein-protein networks.** Network maps of known interactions between proteins coded by the PLS1+ and PLS1− gene sets.

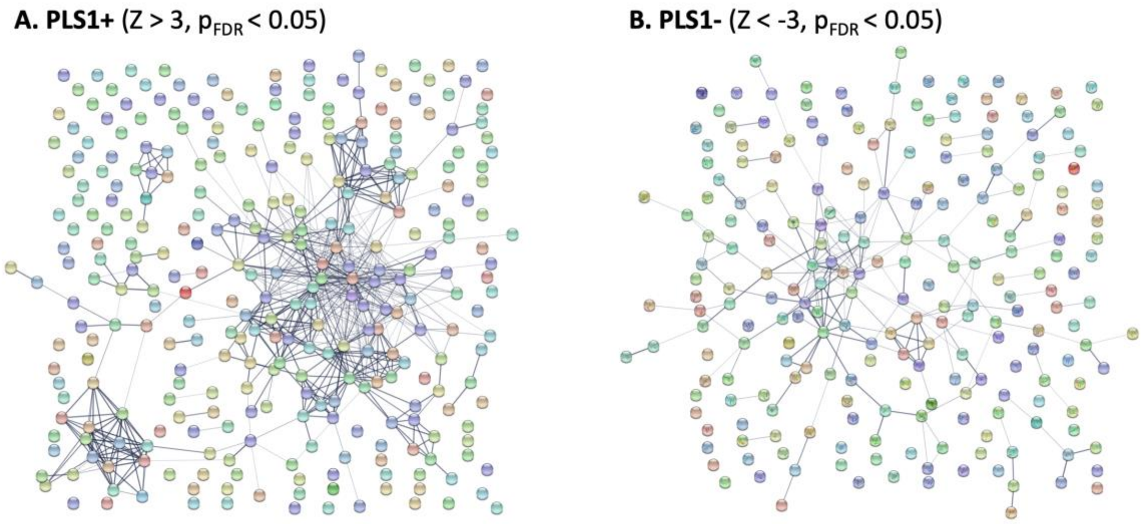


**Supplementary Table S2. Cell-type enrichment analyses (full statistics).** In this table, we present the full statistics of our cell-type enrichment analyses of PLS1+ and PLS1− gene sets. We show odds ratio (OR) for gene enrichment and the respective p-values (before and after FDR correction for the total number of cell types examined).

| **Cell class** | **PLS1+** | | | **PLS1-** | | |
|---|---|---|---|---|---|---|
| | **OR** | **p** | **$p_{FDR}$** | **OR** | **p** | **$p_{FDR}$** |
| **Astrocytes** | **1.24** | **0.02** | **0.04** | 0.21 | 0.99 | 0.99 |
| **Endothelial** | 0.84 | 0.82 | 0.99 | 0.45 | 0.99 | 0.99 |
| **Microglia** | **5.03** | **$4.01 \times 10^{-28}$** | **$2.81 \times 10^{-27}$** | 0.47 | 0.99 | 0.99 |
| **Excitatory Neurons** | 0.23 | 0.99 | 0.99 | **6.06** | **$1.01 \times 10^{-29}$** | **$7.07 \times 10^{-29}$** |
| **Inhibitory Neurons** | 0.18 | 0.99 | 0.99 | **2.46** | **$1.41 \times 10^{-5}$** | **$4.94 \times 10^{-5}$** |
| **Oligodendrocytes** | 0.36 | 0.99 | 0.99 | 0.20 | 0.99 | 0.99 |
| **Oligodendrocyte Progenitor Cells** | **2.14** | **0.001** | **0.004** | 0.73 | 0.76 | 0.99 |

**Supplementary Table S3. Disease-type enrichment analyses (full statistics).** In this table, we present the full statistics of our disease-type enrichment analyses of PLS1+ and PLS1− gene sets. We show odds ratio (OR) for gene enrichment and the respective p-values (before and after FDR correction for the total number of disease types examined).

| **Brain disorder** | **PLS1+** | **PLS1-** |
|---|---|---|

| | OR | p | $p_{FDR}$ | OR | p | $p_{FDR}$ |
|---|---|---|---|---|---|---|
| **Alzheimer** | 1.09 | 0.26 | 0.98 | 0.91 | 0.73 | 0.73 |
| **Parkinson** | 1.05 | 0.40 | 0.98 | 1.13 | 0.28 | 0.63 |
| **Huntington** | 0.99 | 0.55 | 0.98 | 1.09 | 0.41 | 0.63 |
| **Epilepsy** | 0.62 | 0.98 | 0.98 | **1.54** | **0.02** | **0.09** |
| **Autism Sepctrum Disorder** | 0.99 | 0.55 | 0.98 | 0.91 | 0.73 | 0.73 |
| **Major depressive disorder** | 0.76 | 0.90 | 0.98 | **1.57** | **0.02** | **0.09** |
| **Anxiety** | 0.82 | 0.82 | 0.98 | 1.08 | 0.42 | 0.63 |
| **Bipolar disorder** | 0.89 | 0.74 | 0.98 | 0.88 | 0.72 | 0.73 |
| **Schizophrenia** | 0.92 | 0.75 | 0.98 | 1.16 | 0.20 | 0.60 |
